# Activation of plant immunity through conversion of a helper NLR homodimer into a resistosome

**DOI:** 10.1101/2023.12.17.572070

**Authors:** Muniyandi Selvaraj, AmirAli Toghani, Hsuan Pai, Yu Sugihara, Jiorgos Kourelis, Enoch Lok Him Yuen, Tarhan Ibrahim, He Zhao, Rongrong Xie, Abbas Maqbool, Juan Carlos De la Concepcion, Mark J. Banfield, Lida Derevnina, Benjamin Petre, David M. Lawson, Tolga O. Bozkurt, Chih-Hang Wu, Sophien Kamoun, Mauricio P. Contreras

**Affiliations:** The Sainsbury Laboratory, University of East Anglia; Norwich Research Park, Norwich, UK; Imperial College London, London, UK; Department of Biochemistry and Metabolism, John Innes Centre, Norwich, UK

## Abstract

Nucleotide-binding domain and leucine-rich repeat (NLR) proteins can engage in complex interactions to detect pathogens and execute a robust immune response via downstream helper NLRs. However, the biochemical mechanisms of helper NLR activation by upstream sensor NLRs remain poorly understood. Here, we show that the coiled-coil helper NLR NRC2 accumulates *in vivo* as a homodimer that converts into a higher order oligomer upon activation by its upstream virus disease resistance protein Rx. The Cryo-EM structure of NRC2 in its resting state revealed intermolecular interactions that mediate homodimer formation. These dimerization interfaces have diverged between paralogous NRC proteins to insulate critical network nodes and enable redundant immune pathways. Our results expand the molecular mechanisms of NLR activation pointing to transition from homodimers to higher-order oligomeric resistosomes.

## Main Text

The nucleotide-binding and leucine-rich repeat (NLR) class of intracellular immune receptors are an important component of innate immunity across the tree of life. They mediate intracellular recognition of pathogens and subsequently initiate an array of immune responses in order to counteract infection (*1, 2*). Plant NLRs can be activated by pathogen-secreted virulence proteins, termed effectors, which pathogens deliver into host cells to modulate host physiology (*3*). NLRs belong to the signal ATPases with numerous domains (STAND) superfamily. They typically exhibit a tripartite domain architecture consisting of an N-terminal signaling domain, a central nucleotide-binding domain and C-terminal superstructure forming repeats (*4*). The central domain, termed NB-ARC (nucleotide-binding adaptor shared by APAF-1, plant R proteins and CED-4) in plant NLRs, is a signature feature of this protein family and plays a key role as a molecular switch, mediating conformational changes required for activation (*4*). A hallmark of NLR activation in eukaryotes and prokaryotes is their oligomerization into higher-order immune complexes, such as plant resistosomes or mammalian and bacterial inflammasomes. These complexes initiate immune signaling via diverse mechanisms, often leading to a form of programmed cell death (*5–11*). NLRs belong to different phylogenetic clades carrying distinct N-terminal domains with the coiled-coil (CC)-type being the most widespread in plants (*12*). A canonical plant CC-NLR is Arabidopsis ZAR1, which is sequestered in an inactive monomeric state in complex with its partner pseudokinase RKS1. Perception of its cognate effector AvrAC primes individual ZAR1 monomers, which undergo conformational changes and subsequently assemble into pentameric resistosomes (*9, 13*). Following resistosome assembly, CC-NLRs oligomers insert themselves into the plasma membrane and presumably act as calcium channels, initiating immune signaling and cell death, as shown for ZAR1 and the wheat CC-NLR Sr35 (*7, 14*). Oligomerization and calcium channel activity have also been shown for the helper NLR NRG1 (*15, 16*). However, the extent to which the model of plant NLR activation drawn from CC-NLRs like ZAR1 or Sr35 applies to other CC-NLRs remains unknown.

NLR proteins have evolved from multifunctional singletons that can mediate both pathogen effector perception and subsequent immune signaling, as is the case for ZAR1 (*17, 18*). However, several NLRs have sub-functionalized throughout evolution into sensors that are dedicated to pathogen perception and helpers that initiate immune signaling. NLR sensors and helpers can function together as receptor pairs or in higher order configurations termed immune receptor networks (*17*). In the NRC immune receptor network of solanaceous plants, multiple sensor CC-NLRs can signal redundantly via an array of downstream paralogous helper CC-NLRs termed NRCs (NLRs required for cell death) to confer disease resistance to a multitude of pathogens and pests (*19*). Effector perception by NRC-dependent sensors leads to conformational changes which allow their central nucleotide binding (NB) domain to signal to downstream helper NRCs and trigger their oligomerization into a helper resistosome (*20–22*). Although assembly into a resistosome is a shared endpoint between ZAR1 and NRCs, the early stages of NRC activation are not understood. Whether the resting form of NRCs exhibits a similar conformation as the monomeric resting state ZAR1 is not known.

In this work, we used Cryo-EM to solve the structure of the helper NLR NbNRC2 purified from the model plant *Nicotiana benthamiana*. We show that unlike the canonical CC-NLR ZAR1, NbNRC2 forms a homodimer in its resting state. We leveraged our recent finding that the NB domains of sensor NLRs are sufficient to activate downstream NRC helpers (*20*) to study the activation of the NbNRC2 homodimer by the NB domain of the *Potato virus X* disease resistance protein Rx (Rx^NB^). Using *in vitro* reconstitution assays, we show that Rx^NB^ can trigger assembly of a high molecular weight NbNRC2 oligomer. Remarkably, even though NbNRC2, NbNRC3 and NbNRC4 are genetically redundant paralogs for Rx-mediated signaling and disease resistance (*19*), NbNRC2 does not hetero-associate or co-localize with other NRC paralogs from *N. benthamiana*. These findings were corroborated with phylogenomic analysis of the homodimerization interface across the 15 major clades of NRC proteins found in the Solanaceae family. This interface has diverged within the majority of the individual NRC clades likely leading to molecular insulation of helper nodes in the immune receptor network. We propose that insulation of NRC helper pathways emerged throughout evolution to generate genetic redundancy, minimize undesired cross-activation and evade pathogen suppression of immunity. Our work expands our understanding of the mechanisms of NLR activation, pointing to helper NLR conformational transitions from homodimers to oligomeric resistosomes.

## Cryo-EM structure of the NRC2 homodimer

Blue native polyacrylamide gel electrophoresis (BN-PAGE) indicates that the helper NLR NbNRC2 accumulates *in planta* at a higher molecular weight than a monomer (*20–23*). This prompted us to determine whether this protein self-associates *in planta* in its inactive state. Co-immunoprecipitation assays conducted in the solanaceous model plant *N. benthamiana* revealed that NbNRC2 self-associates unlike the canonical functional singleton ZAR1 (**Fig. 1A**).

**Fig. 1:**
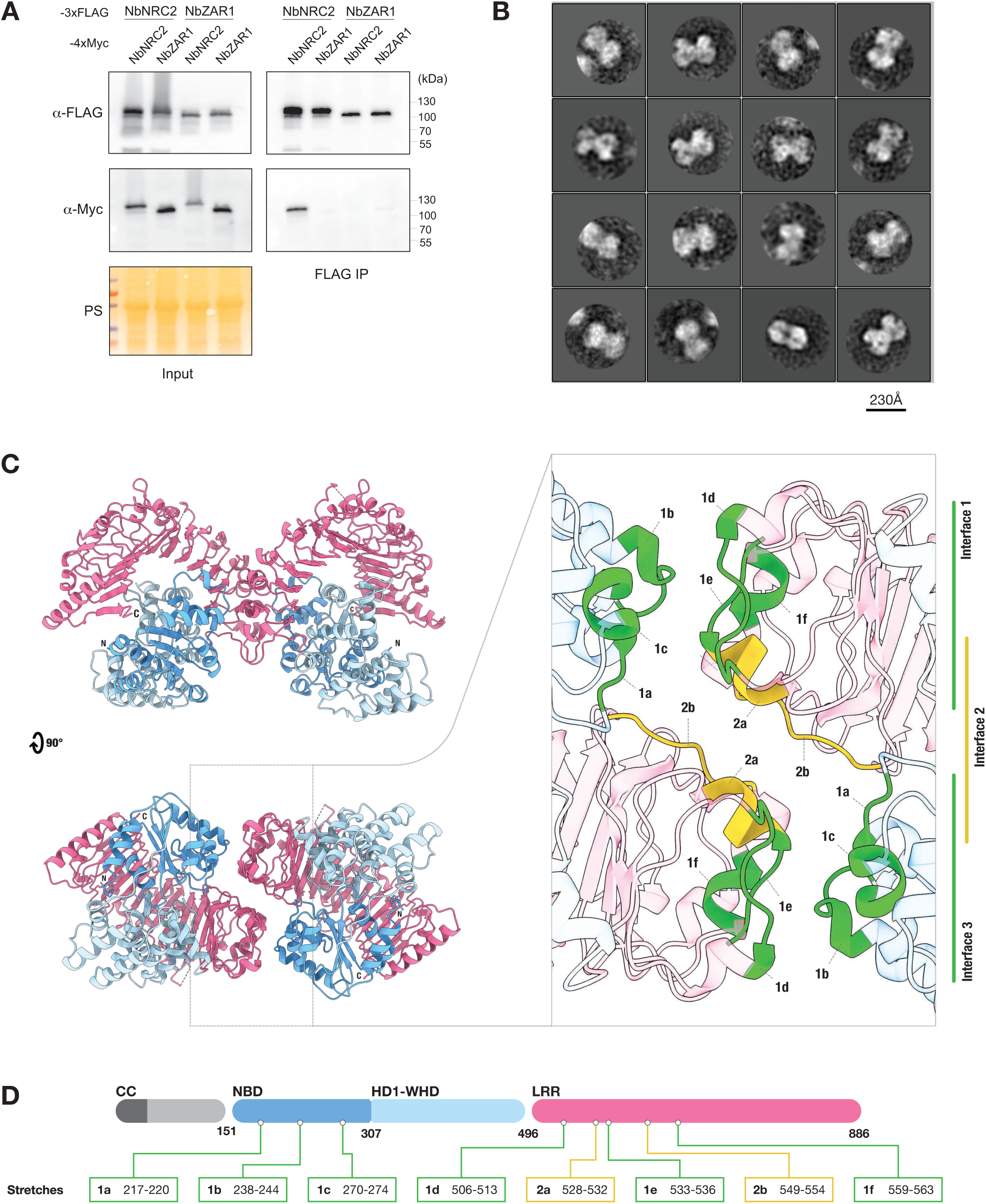
The helper NLR NRC2 forms a homodimer *in planta*. (**A**) Co-immunoprecipitation assays to test self-association of NRC2 and ZAR1 from *N. benthamiana*. C-terminally 3xFLAG-tagged NbNRC2 and NbZAR1 proteins were coexpressed with C-terminally 4xMyc tagged NbNRC2 or NbZAR1. Immunoprecipitations were performed with agarose beads conjugated to FLAG antibodies (FLAG IP). Total protein extracts were immunoblotted with the antisera labelled on the left. Approximate molecular weights (in kDa) of the proteins are shown on the right. Protein loading control was carried out using Ponceau stain (PS). The experiment was performed three times with similar results. (**B**) 2D classifications from negative staining transmission electron microscopy images of affinity-purified NbNRC2. NRC2 appears as a dimer, with a maximum dimension of approximately 150 Å at its widest point. Scale bar = 230 Å. (**C**) Cryo-EM structure of resting state of NRC2 dimers. Atomic model corresponding to NRC2 homodimer shown in two orthogonal views, with resolved NB domain (NBD), HD1-WHD and LRR domains. Notably, the N-terminal CC domain of NRC2 was absent from the Cryo-EM density. Inset shows details of interfaces between the two NRC2 protomers, highlighting amino acid stretches corresponding to three contact interfaces. (**D**) Color coding of the domains is shown in the schematic representation of the domain architecture and boundaries of NRC2, which includes the exact boundaries of the amino acid stretches at the dimerization interfaces. A schematic representation of the pipeline used for Cryo-EM imaging, data processing and model building can be found in **Fig. S2**. Additional views of the structure and dimerization interface can be found in **Fig. S3, Fig. S4 and Movie S1**. Additional information on image processing and model building can be found in **Table S1**.

To further investigate the inactive form of NbNRC2, we transiently overexpressed it in leaves of *N. benthamiana*. Analytical gel filtration assays using affinity-purified NbNRC2 protein revealed elution volumes consistent with a helper NLR homodimer (**Fig. S1**). Subsequent negative-staining imaging of purified NbNRC2 and their class averages also suggested an NRC2 homodimer (**Fig. 1B**). Cryo-EM imaging of these particles and single particle reconstruction yielded a 3.9 Å average resolution structure of the homodimer with C2 symmetry (**Fig. 1C, Fig. S2, Fig. S3, Fig. S4, Table S1**). The reconstructed structure contains the NB-ARC and LRR domains of NbNRC2, with the N-terminal CC domain appearing disordered in the density map. Flexible fitting of the AlphaFold2 model of NbNRC2 monomers into the Cryo-EM map of the dimeric assembly revealed the relative orientation of the helper NLR protomers as well as the domains responsible for homodimer organization (**Fig. 1C**, **Fig. 1D, Fig. S3, Fig. S4, Movie S1**).

The dimerization interface can be divided into 3 parts. The NB domain of each NbNRC2 protomer interacts with the LRR region of the other protomer, forming two similar interfaces related by 2-fold symmetry of the dimer (interface 1 and 3) (**Fig. 1C**). The N-terminal LRR region of each NbNRC2 protomer stacks over the equivalent region of the other protomer, at the two-fold symmetry, forming interface 2 (**Fig. 1C**). These three interfaces hold the homodimer together with a buried surface area 937 Å^2^ for each protomer. The residues at the dimerization interface were listed with a distance cut-off of 6 Å (**Table S2**). Interfaces 1 and 3 are composed of three stretches of residues in the NB domain (Stretch 1a-1c, residues 217-220, 238-244 and 270-274 respectively) in one protomer and three stretches of residues in the LRR (stretches 1d-1f, residues 506-513, 533-536 and 559-563 respectively) from the other protomer (**Fig. 1C**, **Fig. 1D**). Two stretches of residues in the LRR (stretches 2a and 2b, residues 528-532 and 549-554, respectively) stack together forming interface 2 (**Fig. 1C**, **Fig. 1D**). In the dimer assembly, the C-termini of the LRR domain and the NB domain of each protomer interact *in cis*. The 15 parallel strands of each LRR domain orient antiparallel relative to the other protomer in the dimer. The C-termini of each LRR are found at the periphery of the assembly and do no not contribute to the dimerization interface (**Fig. 1C**).

## NRC2 homodimer converts into a resistosome upon activation

To further study NRC2 activation, we set up *in vitro* reconstitution experiments with purified NbNRC2 and carried out BN-PAGE assays. First, to determine the optimal experimental conditions, we incubated purified NbNRC2 with increasing levels of ATP, which is necessary for NLR activation (*4, 24*). Incubation of NbNRC2 with 15 mM ATP led to the disappearance of the lower molecular weight NRC2 band and appearance of higher molecular weight complexes migrating at a similar size to the NbNRC2 oligomer observed *in planta* (*20–23*) (**Fig. 2A**). This indicates that purified NbNRC2 may get spontaneously activated at 15 mM ATP *in vitro*. We selected 5 mM ATP for subsequent assays.

**Fig. 2:**
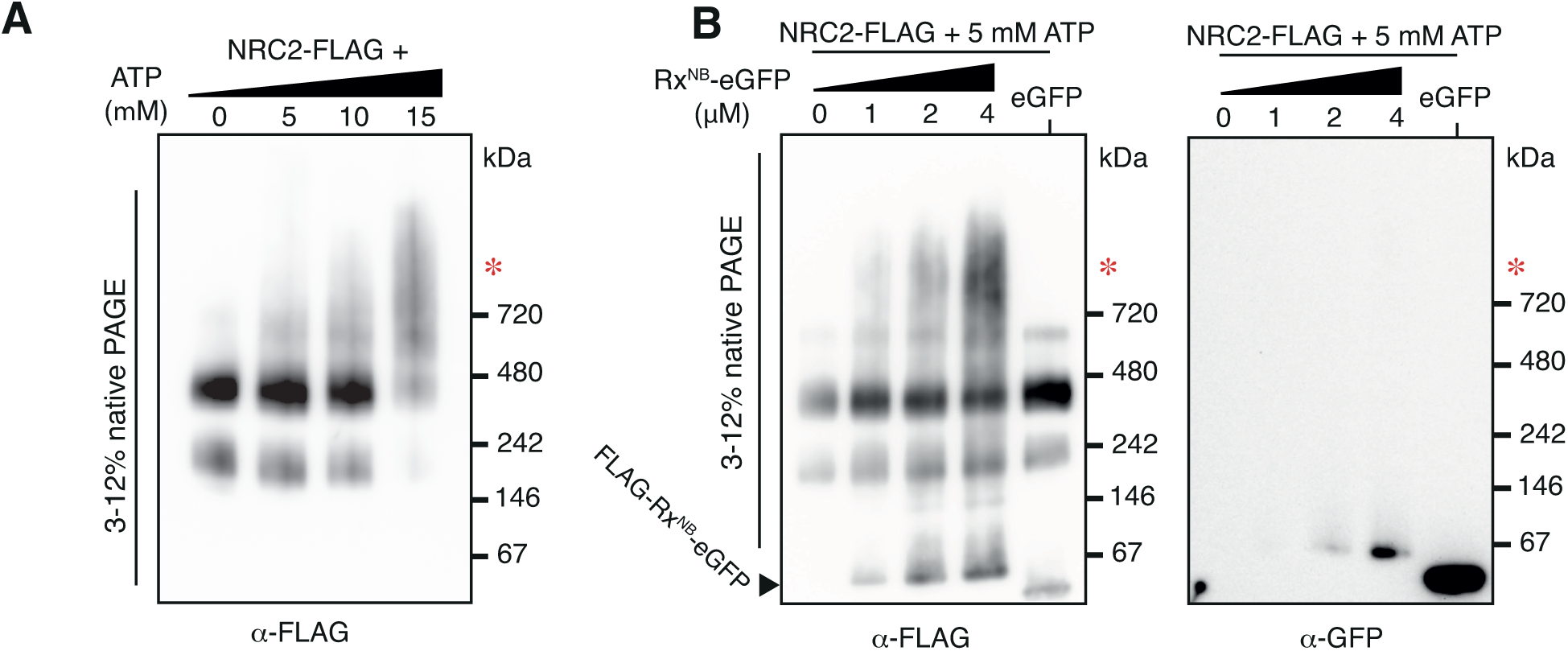
The nucleotide-binding domain of the sensor NLR Rx converts the NRC2 helper homodimer into a higher order oligomer *in vitro*. (**A**): BN-PAGE assay with NRC2 incubated with ATP *in vitro*. Purified NbNRC2-FLAG (5 μM) was incubated overnight with no ATP or with increasing amounts of ATP (5, 10 or 15 mM). The experiment was repeated three times with similar results. (**B**) BN-PAGE assay with NRC2 incubated with Rx^NB^ *in vitro*. Purified NbNRC2-FLAG (5 μM) was incubated overnight with increasing amounts of purified FLAG-Rx^NB^-eGFP (0, 1, 2 and 4 μM, respectively). eGFP (4 μM) was included as a negative control. The experiment was performed three times with similar results. (**A-B**) *In vitro* reactions were run in parallel on native and denaturing PAGE and immunoblotted with the antisera labelled below each blot. SDS-PAGE blots accompanying BN-PAGE assays in panel **B** are found in **Fig. S5**. Approximate molecular weights (in kDa) of the proteins are shown on the right. Red asterisk indicates signal at a molecular weight matching the previously reported NRC2 resistosome-like oligomer.

Next, we hypothesized that the disease resistance protein Rx disrupts the NbNRC2 homodimer to trigger activation of its helper NLR. To test this, we used Rx^NB^, the 154 amino acid NB domain of Rx, which is necessary and sufficient to activate NbNRC2 in the absence of a pathogen effector (*20, 25*). In the presence of 5 mM ATP, these *in vitro* reconstitution experiments revealed that increasing amounts of Rx^NB^ triggered the appearance of higher order complexes similar in size to the NbNRC2 resistosome (**Fig. 2B, Fig. S5**). We conclude that conversion of the helper NLR homodimer into a resistosome by a sensor NLR is a key step of NbNRC2 activation in a mechanistic model that is different from the activation of the ZAR1/RKS1 complex (*9, 13*).

## NRC paralogs do not associate with each other

Paralogous NRC helpers can form genetically redundant network nodes in the ∼100-million-year-old immune receptor network, therefore promoting robustness and evolvability of the plant immune system (*3, 19, 26*). However, the biochemical basis of NRC redundancy is unknown. We hypothesized that NRC paralogs acquired mutations in their dimerization interfaces that insulate them from one another, as predicted from prior theory on the evolution of orthogonal signaling pathways following gene duplication events (*27, 28*). We investigated the degree to which NRC paralogs form heterodimers with each other using *in planta* co-immunoprecipitation assays of resting forms of NbNRC2, NbNRC3 and NbNRC4, and used NbZAR1 as a negative control. Our assays confirmed NbNRC2 self-association but yielded weak to non-detectable signal for hetero-association between NbNRC2 and NbNRC3, NbNRC4 or NbZAR1 (**Fig. 3A**). We conclude that NbNRC2 preferentially self-associates in *N. benthamiana*, and does not form stable heterocomplexes with its paralogs NbNRC3 and NbNRC4.

**Fig. 3:**
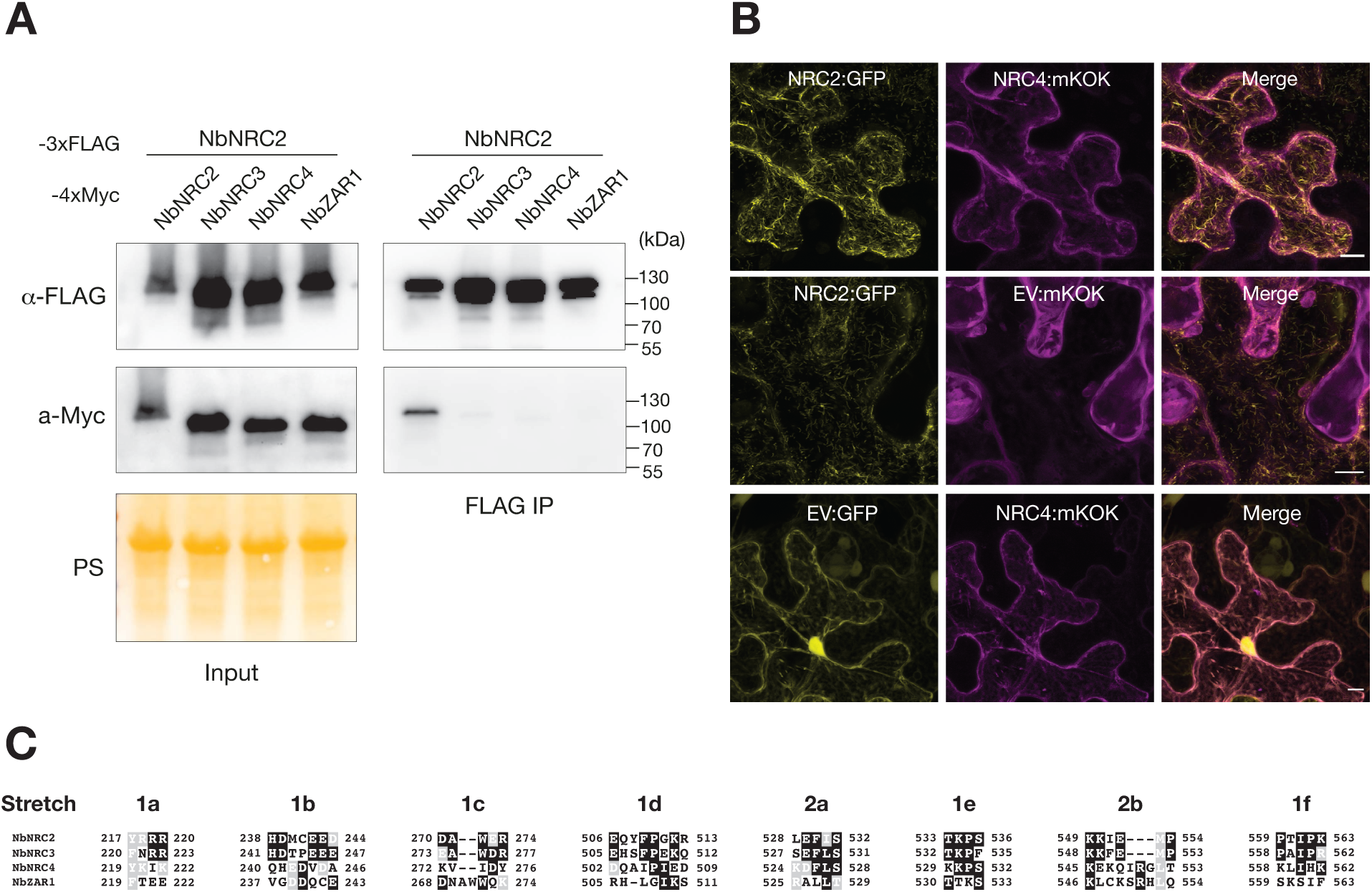
NRC2 does not hetero-associate or co-localize with other NRC paralogs in its resting state. (**A**) Co-IP assays to test the association of NRC2 with its paralogs NRC3 and NRC4 from *N. benthamiana*. C-terminally FLAG-tagged NbNRC2 was co-expressed with C-terminally 4xMyc-tagged NbNRC2, NbNRC3, NbNRC4 or NbZAR1 as a negative control. IPs were performed with agarose beads conjugated to FLAG antibodies (FLAG IP). Total protein extracts were immunoblotted with the antisera labelled on the left. Approximate molecular weights (kDa) of the proteins are shown on the right. Protein loading control was carried out using Ponceau stain (PS). The experiment was performed three times with similar results. (**B**) Z-stacks of confocal micrographs show the localization of NbNRC2:GFP together with NbNRC4:mKOk or EV:mKOk. C-terminally GFP-tagged NbNRC2 was co-infiltrated with either NbNRC4:mKOk or EV:mKOK in wildtype *N. benthamiana* plants. EV:GFP was co-infiltrated with NbNRC4:mKOk as an additional control. Scale bars represent 10 μm. Images were taken 48 hours post agroinfiltration. (**C**) Amino acid sequence alignment of the eight stretches identified at the NbNRC2 dimerization interface, comparing NbNRC2, NbNRC3, NbNRC4 and NbZAR1.

We independently corroborated these findings using live-cell confocal microscopy (**Fig. 3B**). NbNRC2 accumulates in filament-like structures in its resting state that do not overlap with the previously reported peripheral subcellular localization of NbNRC4 (*29*) further supporting the view that these critical network nodes signal via insulated pathways. Overall, these findings are consistent with the distinct sequences of the dimer interfaces of NRC helper NLRs inferred from the NbNRC2 homodimer structure (**Fig. 3C, Data S1**).

## Homodimerization interfaces have diverged throughout NRC evolution

The observation that NbNRC2 does not form heterocomplexes with other NRC paralogs despite their common evolutionary origin prompted us to perform phylogenomic analyses across the highly expanded NRC clade of the Solanaceae family. First, we developed a computational pipeline to extract 1,092 NRC sequences from 6,630,292 predicted proteins from 123 solanaceous genome assemblies representing 38 species of the genera *Solanum*, *Physalis*, *Capsicum* and *Nicotiana* (**Data S2**). We grouped these NRC sequences into 15 phylogenetic clades ranging in size from 31 to 129 sequences (**Fig. S6**). 10 of the 15 clades include orthologs from at least two genera, therefore representing relatively deep NRC lineages (**Fig. S7**).

The consensus sequences of the eight amino acid stretches found in the dimerization interface were, overall, degenerated across all NRC sequences compared to known conserved motifs of CC-NLRs, e.g. P-loop and MHD motifs, suggesting that they have diversified throughout NRC evolution (**Fig. 4A**, **Fig. S7**, **Data S3**). However, the dimerization interfaces tended to be conserved within each clade (**Fig. 4B**, **Fig. S7, Fig. S8, Data S4**). We identified unique polymorphisms within the major NRC2, NRC3, and NRC4 clades by calculating the relative amino acid ratio for one clade vs. the others using a fifty-fold cutoff with a minimum frequency of 80% within the query clade (**Fig. 4B**). Using these criteria, the NRC2 and NRC3 clades, which contain representative proteins used in co-immunoprecipitation assays in **Fig 3A**, had 5 and 3 unique polymorphisms among the 44 amino acids that map to the homodimer interface (**Fig. 4B**, **Data S5**).

**Fig. 4:**
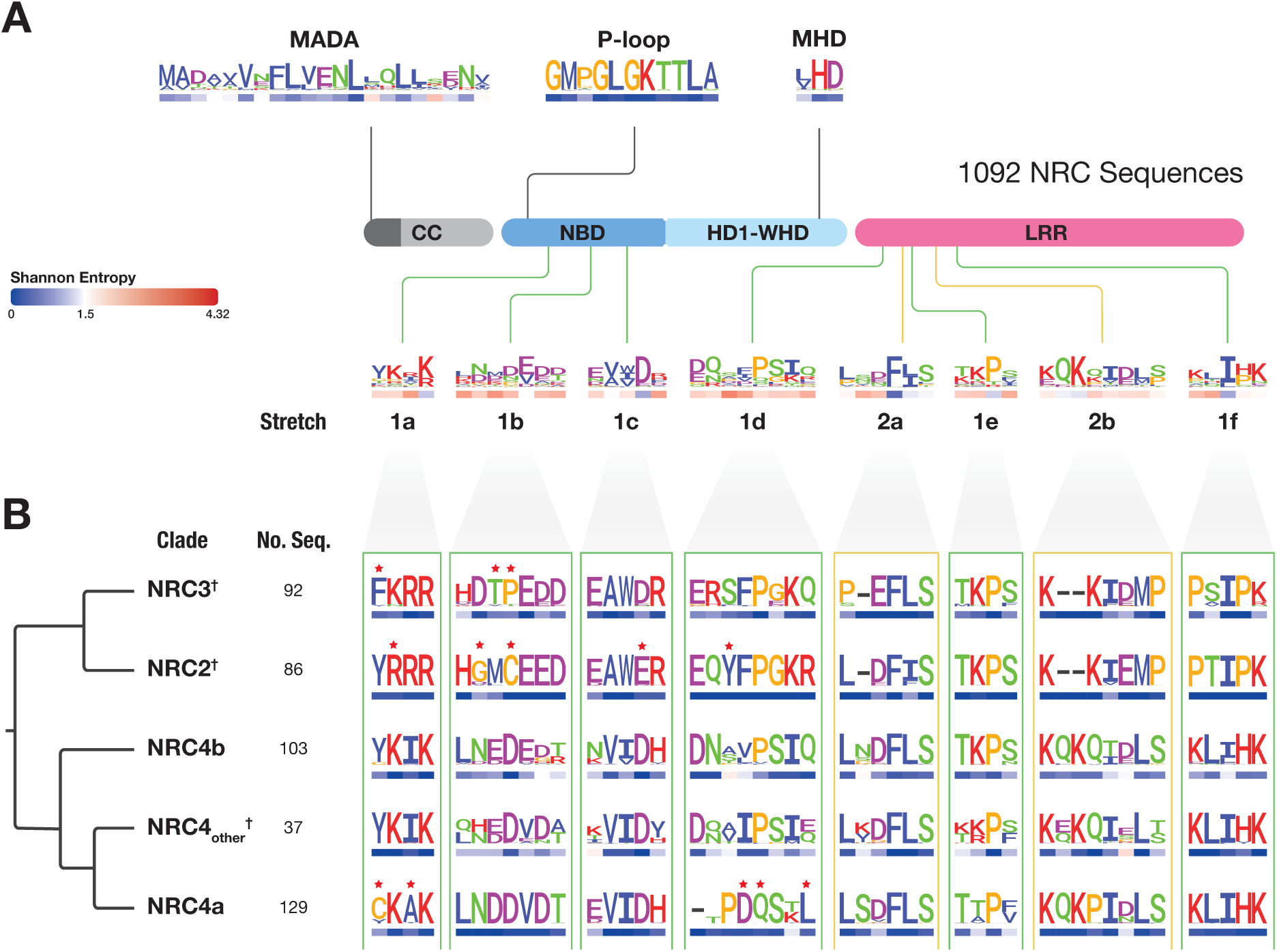
The NRC homodimerization interface has diversified throughout evolution of NRC clade. (**A**) Schematic representation of NRC domain architecture, featuring CC, NBD, HD1-WHD, and LRR domains. Approximate position of different previously characterized NLR motifs (MADA, p-loop, MHD) and the eight amino acid stretches present at the dimerization interface are indicated. A database of 993 unique NRC sequences and 99 reference NRC sequences was compiled from a total of 123 solanaceous genomes (**Data S2**) and used to generate consensus sequence patterns for each motif/region (**Data S3**). Shannon entropy scores for each amino acid position were calculated using this database and indicated, in bits, below each position in the consensus sequence pattern (**Data S4**). A Shannon entropy score of 1.5 was established as a threshold to distinguish between highly variable (>1.5 bits) and non-highly variable (<1.5 bits) amino acids, as established previously (*40*). (**B**) All 1,092 NRC sequences were grouped by phylogeny (**Fig. S6**), and consensus sequence patterns within each clade were generated with sequence logos for each of the eight amino acid stretches that define the dimerization interface (**Fig. S7**). NRC clades are ordered by phylogenetic relationship, indicated on the left along with the number of NRC sequences within each clade. Shannon entropy score is indicated below each amino acid in the consensus sequence pattern. Red stars above indicate amino acid polymorphisms that are unique to each clade, as determined by analysing their relative amino acid ratio among NRC clades (**Data S5**). †Indicates clades containing *N. benthamiana* NRCs used in the co-immunoprecipitation experiments in Fig. 3A. More detailed Shannon entropy plots calculated across all 1092 NRC sequences and within each individual NRC clade are shown in **Fig S8**.

Next, we corroborated these findings using Shannon’s entropy analyses. The entropy of a given amino acid position ranges from zero (Log_2_1) for fully conserved residues to a maximum of 4.32 (Log_2_20) when all 20 amino acids are equally found (30). We used a threshold of 1.5 to highlight highly variable positions consistent with previous NLR studies (31). All eight dimerization stretches scored above 1.5 when all 1,092 NRC helpers were compared but were highly conserved within 11 of the 15 NRC clades (**Fig. 4A**, **Fig. S7**, **Fig. S8**). In 6 clades (NRC0, NRCX, NRC2, NRC3, NRC4a and NRC4t), all residues across the eight stretches scored under the 1.5 threshold and showed little to no sequence diversity (**Fig. S7**, **Fig. S8**, **Data S4**). In 5 clades (NRC1, NRC6, NRC7, NRC4b and NRC4_other_), the eight stretches were under the 1.5 threshold except for one to three variable residues within each clade (**Fig. S7**). In the remainder of the clades (NRC8, NRC9, NRC5 and NRC4c), all eight stretches had up to 12 variable residues within each clade with entropy values higher than 1.5 indicating higher levels of sequence diversity compared to other clades (**Data S4**). In contrast, conserved motifs, such as P-loop and MHD, were consistently under the defined threshold with 13 of the 15 clades scoring under 1.5 and displaying little to no sequence variation (**Fig. 4**, **Fig. S7**, **Fig. S8**, **Data S4**).

## Discussion: Contrasting activation mechanisms across plant NLRs

Our study shows that a CC-NLR helper in the NRC immune network exhibits a distinct mechanism of activation relative to the canonical ZAR1/ZRK system (**Fig. 5**). Our data is consistent with a model in which the NB domain of the disease resistance protein Rx converts the homodimer of its downstream helper NbNRC2 into a high molecular weight oligomer. The Cryo-EM structure of the NbNRC2 homodimer revealed novel intermolecular interactions that hold this helper NLR in its resting state. The surfaces involved in NbNRC2 homodimerization are predicted to be surface exposed in the activated NbNRC2 oligomer based on ZAR1 resistosome structure. This implies that conversion from homodimer to a multimeric resistosome is likely to involve dimer dissociation into monomers and additional intermediate assemblies. The contrasting models in early steps of activation of NRC2 and ZAR1 indicate that plant NLRs have evolved a diversity of activation mechanisms possibly associated with their evolutionary transitions from functional singletons to paired and networked immune receptors. Indeed, ZAR1 is more likely to reflect the ancestral state of CC-NLRs given that it is present across ancient plant lineages, including dicotyledons, monocotyledons and magnoliids and has been proposed to guard ZRK proteins since the Jurassic era (*30, 31*). In contrast, NRC sensor/helper pairing is thought to be a more recent innovation, having emerged in the Caryophyllales and Asterid lineages of dicotyledoneous flowering plants (*19, 32, 33*). It will be interesting to determine the extent to which homodimerization has evolved as a steady state across NLRs. Already, RPM1, a CC-NLR outside the NRC superclade, is known to self-associate in its resting state (*34*).

**Fig. 5:**
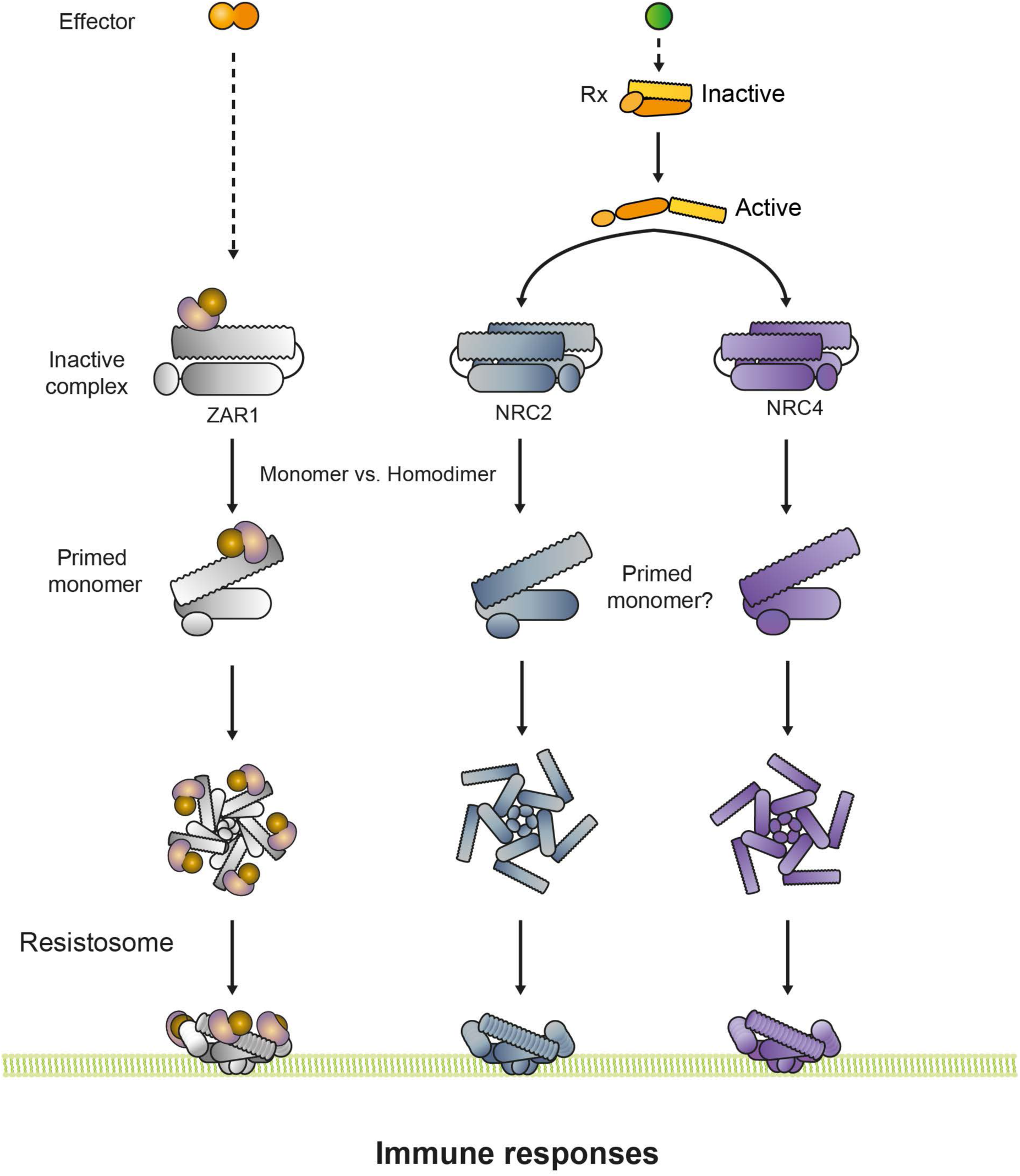
Contrasting mechanisms of activation between ZAR1 and the helper NLR NRC2. The figure depicts the working model for activation of NRC2 homodimers. Unlike the functional singleton ZAR1, the helper NLR NRC2 exists as a homodimer in its resting form. NbNRC2 carries unique polymorphisms at the interface between its two protomers and exclusively associates with itself without hetero-dimerizing with its NRC paralogs NbNRC3 or NbNRC4. Activation of upstream sensors and conditional exposure of their nucleotide binding (NB) domains results in conversion of the NbNRC2 homodimer into a high molecular weight resistosome, presumably via homodimer dissociation into a putative primed monomer which subsequently oligomerizes.

NLRs are generally thought to be hypervariable due to coevolution with rapidly evolving pathogens (*35, 36*). While the NRC clade of helper NLRs exhibits greater conservation within the Solanaceae compared to NRC-dependent sensors and most other NLRs (*19, 32, 33*), these proteins exhibit variable regions across the NRC helper clade (**Fig. 4**, **Fig. S7, Fig. S8**). Our analyses revealed that distinct NRC clades have diverged from each other in the homodimerization interface and that NRC2 does not associate with its NRC3 and NRC4 paralogs. We propose that following duplication, NRC paralogs have accumulated mutations in the dimerization interface that prevent them from forming heterodimers, thereby forming insulated pathways in the immune network. Indeed, redundant nodes in immune receptor networks may help the plant evade suppression by pathogen effectors, contributing to the robustness of the immune system (**Fig. 5**) (*18, 37, 38*).

## Acknowledgments

We are very thankful to D. Lüdke (The Sainsbury Laboratory, Norwich, UK) for valuable discussions and critical reading of this manuscript. We thank C. McClune, (Stanford, California, USA) for useful discussions. We thank J. Richardson (John Innes Centre, Norwich, UK) for support and excellent maintenance of John Innes Centre Bioimaging facility. Negative stain EM data for this investigation were collected at JIC Bioimaging facility, which is supported by the Biotechnology and Biological Sciences Research Council (BB/CCG2240/1). We thank N. Lukoyanova (Birkbeck, University of London, London, UK) for support with Cryo-EM imaging. Cryo-EM data for this investigation were collected at ISMB EM facility, which is supported by the Wellcome Trust (202679/Z/16/Z and 206166/Z/17/Z). S.K. and M.P.C. thank F. Sinatra and L. Messi for inspiration. We thank all members of the TSL Support Services for their invaluable assistance.

## Funding

The authors received funding from the sources listed below. The funders had no role in the study design, data collection and analysis, decision to publish, or preparation of the manuscript.

The Gatsby Charitable Foundation (MS, AT, HP, YS, JK, HZ, RX, AM, BP, LD, DML, CHW, SK, MPC)

The John Innes Foundation (JCDLC)

Biotechnology and Biological Sciences Research Council (BBSRC) BB/P012574 (Plant Health ISP) (MS, AT, HP, YS, JK, HZ, RX, AM, BP, LD, DML, CHW, SK, MPC)

BBSRC BBS/E/J/000PR9795 (Plant Health ISP – Recognition) (MS, AT, HP, YS, JK, RX, AM, BP, LD, DML, CHW, SK, MPC)

BBSRC BBS/E/J/000PR9796 (Plant Health ISP – Response) (MS, AT, HP, YS, JK, RX, AM, BP, LD, DML, CHW, SK, MPC)

BBSRC BBS/E/J/000PR9797 (Plant Health ISP – Susceptibility) (MS, AT, HP, YS, JK, RX, AM, BP, LD, DML, CHW, SK, MPC)

BBSRC BBS/E/J/000PR9798 (Plant Health ISP – Evolution) (MS, AT, HP, YS, JK, RX, AM, BP, LD, DML, CHW, SK, MPC)

BBSRC BB/V002937/1 (LD, SK, MPC) BBSRC BB/T006102/1 (TI, TOB)

BBSRC impact acceleration award BB/X511055/1 (ELHY)

Academia Sinica and National Science and Technology Council NSTC-111-2628-B-001-023 (CHW)

European Research Council (ERC) 743165 (SK)

## Author contributions

Conceptualization: MS, TOB, CHW, SK, MPC

Methodology: MS, AT, HP, YS, JK, ELHY, HZ, RX, AM, JCDLC, BP, MPC

Investigation: MS, AT, HP, YS, JK, ELHY, TI, RX, AM, JCDLC, LD, DML, CHW, MPC

Visualization: MS, AT, HP, ELHY, MPC

Funding acquisition: TOB, SK

Project administration: MS, SK, MPC

Supervision: MJB, SK, MPC

Writing – original draft: MS, AT, YS, SK, MPC

Writing – review & editing: MS, AT, HP, YS, JK, ELHY, TI, HZ, AM, JCDLC, MJB, LD, BP, DML, CHW, TOB, SK, MPC

## Competing interests

TOB and SK receive funding from industry on NLR biology and cofounded a start-up company (Resurrect Bio Ltd.) on resurrecting disease resistance. JK, LD, SK and MPC have filed patents on NLR biology. LD and MPC have received fees from Resurrect Bio Ltd.

## Data and materials availability

All data are available in the main text or the supplementary materials. Data related to the NRC2 structure can be found at PDB/EMDB, with PDB ID 8RFH and EMD-19121, as well as **Table S1**. Datasets associated to NRC phylogenomics analyses can be accessed on Zenodo (*39*).

## Supplementary Materials

Materials and Methods

Figs. S1 to S8

Table S1 to S2

Data S1 to S10

References (41 to 61)

## Supplementary Materials for

### Materials and Methods

#### Plant growth conditions

Wild-type and *nrc2/3/4* CRISPR mutant (*41*) *Nicotiana benthamiana* lines were grown in a controlled environment growth chamber with a temperature range of 22 to 25 °C, humidity of 45% to 65% and a 16/8-hour light/dark cycle.

#### Plasmid construction

We used the Golden Gate Modular Cloning (MoClo) kit (*42*) and the MoClo plants part kit (*43*) for cloning. All vectors used were generated with these kits unless otherwise stated. Cloning design and sequence analysis were done using Geneious Prime (v2021.2.2; https://www.geneious.com). All NRC and ZAR1 constructs used for co-immunoprecipitation assays were cloned into the pICH86988 level one acceptor with a C-terminal 3xFLAG tag (pICSL5007) or a C-terminal 4xMyc tag (pICSL5010). Constructs for protein purification were cloned into pJK268c-mRFP acceptor with 2×35s promoter (pICSL51288) and terminator (pICSL41414). NbNRC2 was cloned with a C-terminal 6xHis-3xFLAG tag. Rx^NB^-eGFP and eGFP constructs were cloned with an N-terminal 6xHis-3xFLAG tag. Constructs used for cell biology were previously reported (*21, 29*).

#### Transient protein expression by agroinfiltration in *Nicotiana benthamiana*

Proteins of interest were transiently expressed in *N. benthamiana* according to previously described methods (*21*). Briefly, leaves from 4–5-week-old plants were infiltrated with suspensions of *Agrobacterium tumefaciens* GV3101 pM90 strains transformed with expression vectors coding for different proteins indicated. Final OD_600_ of all *A. tumefaciens* suspensions were adjusted in infiltration buffer (10 mM MES, 10 mM MgCl_2,_ and 150 μM acetosyringone (pH 5.6)). Final OD_600_ used for each construct was 0.3.

#### Co-Immunoprecipitation (co-IP) assays

Co-Immunoprecipitation assays were performed as described previously (*37*). Four to five-week-old *N. benthamiana* plants were agroinfiltrated as described above with constructs of interest and leaf tissue was collected 3 days post agroinfiltration. Final OD_600_ used for each construct was 0.3. Leaf tissue was ground using a Geno/Grinder tissue homogenizer. GTEN extraction buffer was used (10% glycerol, 50 mM Tris-HCl (pH 7.5), 5 mM MgCl_2_ and 50 mM NaCl) supplemented with 10 mM DTT, 1x protease inhibitor cocktail (SIGMA) and 0.2% IGEPAL (SIGMA). Samples were incubated in extraction buffer on ice for 10 minutes with short vortex mixing every 2 minutes. Following incubation, samples were centrifuged at 5,000 x*g* for 15 minutes and the supernatant was collected. This was spun down an additional time at 5,000 xg for 15 minutes and then supernatant was filtered using Minisart 0.45 μm filter (Sartorius Stedim Biotech, Goettingen, Germany).

Part of the extract was set aside prior to immunoprecipitation. These were the inputs. 1.4 ml of the remaining filtered total protein extract was mixed with 30 μl of anti-FLAG agarose beads (SIGMA) and incubated end over end for 90 minutes at 4 °C. Beads were washed 5 times with immunoprecipitation wash buffer (GTEN extraction buffer with 0.2% v/v IGEPAL (SIGMA). Associated plant proteins were competitively eluted by excess of 3xFLAG peptides. Elution was spun down at 1000 xg for 1 minute and the supernatant was transferred to a new tube. Inputs and eluted immunoprecipitates were diluted in SDS loading dye and denatured by heating for 10 minutes at 72 °C. All samples were used for SDS-PAGE. Briefly, they were run on 4-20% Bio-Rad Mini-PROTEAN TGX gels alongside a PageRuler Plus prestained protein ladder (Thermo Scientific). The proteins were then transferred to polyvinylidene difluoride membranes using Trans-Blot Turbo Transfer Buffer using a Trans-Blot Turbo Transfer System (Bio-Rad) as per the manufacturer’s instructions. Membranes were immunoblotted as described below.

#### Immunoblotting and detection of BN-PAGE and SDS-PAGE assays

Blotted membranes were blocked with 5% milk in Tris-buffered saline plus 0.01% Tween 20 (TBS-T) for an hour at room temperature and subsequently incubated with desired antibodies at 4 °C overnight. Antibodies used were anti-GFP (B-2) HRP (Santa Cruz Biotechnology), anti-FLAG (M2) HRP (Sigma) or anti-Myc (9E10) HRP (Roche), as indicated on the figures, all used in a 1:5000 dilution in 5% milk in TBS-T. To visualize proteins, we used Pierce ECL Western (32106, Thermo Fisher Scientific), supplementing with up to 50% SuperSignal West Femto Maximum Sensitivity Substrate (34095, Thermo Fisher Scientific) when necessary. Membrane imaging was carried out with an ImageQuant LAS 4000 or an ImageQuant 800 luminescent imager (GE Healthcare Life Sciences, Piscataway, NJ). Rubisco loading control was stained using Ponceau S (Sigma) or Ponceau 4R (A.G. Barr).

#### Protein purification from *N. benthamiana*

Around 30 leaves of *nrc2/3/4* KO *N. benthamiana* were agroinfiltrated as described above to transiently express NbNRC2-6xHis-3xFLAG, 6xHis-3xFLAG-Rx^NB^-eGFP or 6xHis-3xFLAG-eGFP. All constructs were cloned in pJK268c acceptor. Tissue was harvested after 3 days and snap-frozen in liquid nitrogen. All samples were stored at –80° C until the protein was finally extracted. The entire purification process was completed in the same day for each preparation. On the day of purification, the frozen tissue was ground into fine powder in a mortar and pestle that was pre-cooled with liquid nitrogen. 20 g of ground powder were resuspended with extraction buffer (100 mM Tris-HCl pH 7.5, 150 mM NaCl, 1 mM MgCl_2_ 1 mM EDTA, 10% glycerol, 10 mM DTT, 1x cOmplete EDTA-free protease inhibitor tablets (Sigma) and 0.4% v/v IGEPAL). 20 g of ground tissue was resuspended in ice-cold extraction buffer at a 1:4 w/v ratio (20 g of powder in 80 mL of extraction buffer). After vortexing and resuspending the powder in this buffer, the crude extract was spun down for 10 minutes at max. speed. The supernatant was transferred to a new tube and centrifuged again for 10 minutes at max. speed. This second supernatant was filtered using Miracloth (Merck). 200 μl of anti-FLAG beads were added to the supernatant and incubated at 4° C for 90 minutes. The tubes were in constant rotation to prevent the beads from sedimenting. The protein-bound beads were collected on an open column and washed with 10ml of wash buffer (100 mM Tris-HCl pH 7.5, 150 mM NaCl, 1 mM MgCl_2_ 1 mM EDTA, 10% glycerol and 0.4% v/v IGEPAL). The washed beads were collected in an Eppendorf tube and eluted with the final isolation buffer in 200 μl volume (100mM Tris HCl (pH 7.5), 150mM NaCl, 1mM MgCl2, 1mM EDTA, protease inhibitor cocktail, 5% glycerol and 0.2% IGEPAL) supplemented with 0.3mg/ml 3xFLAG peptide. The eluted protein was used for SDS-PAGE to assess sample quality and purity. About 1.2 to 1.5 mg/ml concentration was obtained for NRC2, Rx^NB^-eGFP and eGFP from each purification, as determined by absorption at 280 nm. FLAG-eluted pure protein samples were used for all invitro experiments and structural studies.

#### Gel filtration assays with purified NbNRC2

C-terminally 6xHis-3xFLAG NbNRC2 was transiently expressed in leaves of nrc2/3/4 KO *N. benthamiana* as described above. C-terminally 6xHis-3xFLAG-tagged NbNRC2 used in gel filtration assays was cloned in pICH86988 acceptor. Extracts were purified as described above and injected onto Superdex 200 Increase 10/300 GL column (GE Healthcare) connected to an AKTA FPLC system (GE Healthcare). The column was pre-equilibrated in buffer containing 25 mM Tris-HCL pH 8, 500mM NaCl, 5% glycerol and 1 mM EDTA. Samples were centrifuged prior to loading. The sample was injected at a flow rate of 0.2 ml/min at 4°C and 0.5 ml fractions were collected for analysis by SDS-PAGE gels.

#### Negative staining

3 μl of purified NRC2 sample diluted to 0.3mg/ml were applied to 400 mesh Copper grids (Agar Scientific) with continuous carbon that was glow discharged using a PELCO Easiglow (Ted Pella) for 30 seconds at 8 mA. After 30 seconds, the sample was blotted using Sartorius 292 filter paper and immediately washed with two consecutive drops of 100 μl distilled water each. After washing, 10 μl of 2% (w/v) uranyl acetate stain in H_2_O was applied. After 20 seconds, the excess was blotted off. Immediately after this, another 10 μl of 2% (w/v) uranyl acetate stain in H_2_0 was applied for 30 seconds and then blotted off. The grid was air-dried and examined in a FEI Talos F200C transmission electron microscope (Thermo Fisher Scientfific), operated at 200 keV, equipped with a 4k OneView CMOS detector (Gatan). 15 representative micrographs were recorded with a magnified pixel size 1.7Å (**Fig. 1B**) each with a high particle density. Particles were auto-picked from these 15 micrographs using RELION 3.1 without any template. 2D-classification of the auto-picked particles (2447 particles) from these images revealed particles resembling putative NRC2 as dimers (**Fig. 1B**).

#### Cryo-EM sample preparation and data collection

Schematic representation of the Cryo-EM structural determination workflow can be found in **Fig. S2**. C-flat grids (Protochips: 1.2/1.3 300 mesh) were used for Cryo-EM grid preparation. 3 μl of 1.2 mg/ml of full-length NRC2 sample were applied two times over negatively glow discharged C-flat grids and coated with graphene oxide. The sample was applied inside the chamber of a Vitrobot Mark IV (Thermo Fisher Scientific) at 4° C and 90% humidity and vitrified in liquid ethane. Without graphene oxide coating, no particles were observed in vitreous ice, so graphene-coated grids were used. The Cryo-EM images were collected on a FEI Titan Krios (Thermo Fisher Scientific) operated at 300 keV equipped with a K3 direct electron detector (Gatan) after inserting an energy filter with a slit width of 20 eV. Micrographs were collected with a total dose of 50 e^−^/Å^2^, nominal magnification of 105 kx, giving a magnified pixel size of 0.828 Å. Images were collected as movies of 50 fractions with defocus values ranging from –1.5 µm to –2.7 µm with 2 exposures per hole. A total of 6135 movies were collected for the image processing and 3D reconstruction using RELION 4.0 (*44*).

The micrograph movies were imported into RELION and subjected to drift correction using MotionCor2 (*45*) and CTFFIND4.0 was used for fitting of contrast transfer function and defocus estimation. The Laplacian-of-Gaussian auto-picker in RELION 4.0 was used for automatic reference-free particle picking (*44*). The particles were extracted in 256-pixel box and subjected to several rounds of 2D classification with a circular mask of 150 Å. The 2D classes revealed a clear dimer with secondary structural features in many views. Clean 2D-classes were selected (664,305 particles) and were subjected to 3D classification using an *ab initio* 3D model generated by RELION. The best 3D class revealing the protein fold consisted of 229,347 particles and was subjected to 3D refinement using Refine3D using that 3D class as a reference with a circular mask of 180 Å. Following examination of the C1 refined map, a C2 symmetry was applied. After particle polishing, CTF refinement and postprocessing, the final average resolution was 3.9 Å as estimated by the Fourier shell correlation (FSC = 0.143). The local resolution plot was calculated using PHENIX 3.0 (*46*). The final map revealed distinct domain boundaries of the NRC2 protein fold with clear secondary structural features, helical twists and individual beta strands and allowed us to place each NRC2 protomer and initiate model building at this resolution. A significant number of particle views are down the short dimension, owing to graphene oxide backing. The final map revealed the NB-ARC and LRR domains of the NRC2 molecule, with the N-terminal CC domain not resolved.

#### Model building and refinement

The AlphaFold2 model of NRC2 monomer was divided into NB, ARC and LRR domains. Each of these domains was fitted separately into the Cryo-EM density using UCSF ChimeraX as rigid body units (*47*). Significantly, AlphaFold2 multimer was unable to predict the quaternary structure we observed experimentally. The model was then manually adjusted in COOT, combined and refined using the PHENIX real space refinement module and the fit of the atomic model to the features in the Cryo-EM map was evaluated visually (**Fig. S3**) (*46, 48*). This model was validated using PHENIX and MolProbity (*46, 49*). Figures were made using ChimeraX. The interface residues and the buried surface area were evaluated using PyMol and the PISA server (*50, 51*). A distance cut-off 6 Å is used to define residues at the interface to accommodate all short– and long–range interacting residues. These residues compose the eight dimerization stretches indicated in **Fig. 1C** and listed in **Table S2**. These are the stretches carried for subsequent phylogenetic analyses. Further details on Cryo-EM data processing statistics can be found in **Table S1**.

#### *In vitro* NRC2 activation assays

To understand the impact of excess ATP on NRC2, 6 μl of 10 μM NbNRC2 were incubated with increasing concentration of ATP (0, 5, 10 and 15 mM respectively) overnight in a 20 μl reaction volume, in presence of 5mM MgCl_2_. The reaction mixture was subjected to BN-PAGE analysis as described below. To test the role of Rx^NB^-eGFP in NRC2 dimer activation *in vitro*, we overexpressed and purified both proteins separately from *nrc2/3/4* KO *N. benthamiana*. eGFP was included as a negative control for Rx^NB^-eGFP. C-terminally FLAG-tagged NbNRC2 and N-terminally FLAG-tagged Rx^NB^-eGFP and eGFP were purified by FLAG affinity purification as described above. 6 μl of 10 μM NbNRC2 were mixed in a final reaction volume of 20 μl with increasing amounts of Rx^NB^-eGFP (1, 2, 4 μM, respectively), in the presence of 5 mM ATP and 5 mM MgCl_2_. Mixtures were incubated on ice overnight and then ran on BN-PAGE as described below.

#### BN-PAGE assays

For BN-PAGE, samples from in vitro studies described below were diluted as per the manufacturer’s instructions by adding NativePAGE 5% G-250 sample additive, 4x Sample Buffer and water. After dilution, samples were loaded and run on Native PAGE 3%-12% Bis-Tris gels alongside either NativeMark unstained protein standard (Invitrogen) or SERVA Native Marker (SERVA). The proteins were then transferred to polyvinylidene difluoride membranes using NuPAGE Transfer Buffer using a Trans-Blot Turbo Transfer System (Bio-Rad) as per the manufacturer’s instructions. Proteins were fixed to the membranes by incubating with 8% acetic acid for 15 minutes, washed with water and left to dry. Membranes were subsequently re-activated with methanol to correctly visualize the unstained native protein marker. Membranes were immunoblotted as described above.

#### Confocal microscopy

Three to four-week-old plants were agroinfiltrated as described above with constructs of interest. The live cell confocal microscopy analyses were conducted two days after agroinfiltration. To image the infiltrated leaf tissue, they were excised using a size 4 cork borer, submerged in dH_2_O and live-mounted on glass slides. The imaging of the abaxial side of the leaf tissue was performed using a Leica STELLARIS 5 inverted confocal microscope equipped with a 63x water immersion objective lens. The laser excitations for GFP and mKOk tags were 488 nm and 514 nm, respectively. To prevent spectral mixing from different fluorescent tags when imaging samples with multiple tags, sequential scanning between lines was applied. All confocal images are single plane images.

#### Bioinformatic and phylogenetic analyses

NLRtracker (*12*) output from 123 genome assemblies representing 38 species in 4 genera of the Solanaceae (nightshade) family (*39, 52*) was imported in R, and 66,451 full-length NLR sequences were extracted. 23,872 NLR sequences with domain architectures of “CNL” and “CN” as determined by NLRtracker were retained. The NB-ARC domain (equivalent to NB domain and HD1-WHD domains in **Fig. 1**) for these NLRs was also extracted from NLRtracker. Subsequently, the NB-ARC sequences of RefPlantNLR (*12*), tomato NRC helpers (*53*), and pepper NRC helpers (*54*) were added to the extracted NB-ARCs. The NB-ARC domain boundaries were retrieved from NLRtracker output, and only sequences with NB-ARC sequences between 300-400 amino acids long were kept and truncated or unusually long NB-ARC sequences were discarded. This led to retaining 19760 sequences. The resulting sequences were aligned using MAFFT v7.505 [–– anysymbol] (*55*). The alignment was then used to generate a phylogenetic tree using FastTree v2.1.11 [–lg] (*56*).

The NRC-network superclade containing 10,665 sequences (including helpers and sensors) was extracted from the CC-NLR phylogenetic tree by selecting the major branch containing both NRC reference helpers and sensors using Dendroscope v3.8.10 [Options > Advanced Options > Extract Subnetwork] (*57*). The NRC superclade sequences were then realigned and used to create a new phylogenetic tree, from which the NRC helper clade was extracted using the same methods. The resulting 1,796 NRC helper sequences, without the reference sequences, were deduplicated using CD-HIT v4.8.1 [–c 1.00] (*58*). The deduplication was carried out to capture the highest diversity without introducing bias into the analysis due to overrepresented sequences.

The resulting sequences were filtered based on length, retaining only sequences between 750 and 950 amino acids long. As a result, 993 sequences were kept. This decision was based on the length distribution within the NRC helper data and the length of the reference NRC sequences. The Geneious prime v2023.2.1 (https://www.geneious.com/) length graph feature was used for this purpose.

The NB-ARC domain from the remaining 993 sequences were combined with length filtered NB-ARCs (300-400 amino acids) of the reference NRC helpers from *N. benthamiana* (16 sequences), tomato (13 sequences), pepper (10 sequences), potato (42 sequences), and RefPlantNLR (14 sequences), resulting in 1,088 sequences in total, and aligned using MAFFT. The alignment was used to construct a new phylogenetic tree of deduplicated NRC helpers and reference sequences using FastTree. Finally, NRC helper subclades were extracted based on the presence of reference NRCs in well-supported major branches. Sequences that fell outside or between major clades were not included in the grouping of NRC helpers into subclades. The identified clades were then extracted using Dendroscope in the same way as before.

The full set of 1,092 NRC sequences consists of full-length sequences of the final set (993 sequences) and the unfiltered reference sequences of *N. benthamiana* (16 sequences), tomato (13 sequences), pepper (14 sequences), potato (42 sequences), and RefPlantNLR (14 sequences). These were also aligned using MAFFT [––localpair] to identify the dimer stretches in all NRC helpers. The alignment of the combined 1,092 sequences was trimmed to retain only important and well-aligned regions. This was accomplished using ClipKIT v2.0.1 [–m gappy] to remove sites with 90% gaps (*59*).

The dimerization stretches, and conserved sequence motifs were mapped onto the alignment of all NRC helpers based on NbNRC2 and NLRtracker and extracted using the R scripts available at [https://github.com/amiralito/NRC2Dimer]. Logo plots for stretches and sequence motifs were generated using the ggseqlogo R package (*60*).

#### Shannon entropy analysis

Shannon Entropy values for all NRC helpers and each NRC clade separately were calculated using the Entropy R package (*61*) and visualized using the R scripts available at [https://github.com/amiralito/NRC2Dimer].

#### Identification of the unique residues in each NRC clade

To identify the unique residues in the NRC clades, we calculated the relative amino acid ratio among the NRC clades NRC2/3/4a/4b/4_other_. We first divided the sequences into query and subject groups. Based on the trimmed alignment of NRC helper, we calculated the amino acid frequency at each position in both query and subject groups. We then divided the amino acid frequency of the query group by that of the subject group at the same position. We only retained positions where one of the amino acid frequencies is more than 80% in the query group, and we regarded positions with a relative amino acid ratio of more than 50 as unique residues in the query clade. If an amino acid in the query group does not exist in the subject group, the ratio was output as infinite. The gaps were also ignored in this analysis. If all the query sequences had a gap at a position, the ratio was output as ‘NA’. We iteratively calculated this relative amino acid ratio between an NRC clade and the other NRC clades, by alternating the query NRC clade. The scripts for this analysis are also available at [https://github.com/amiralito/NRC2Dimer].

**Fig. S1:**
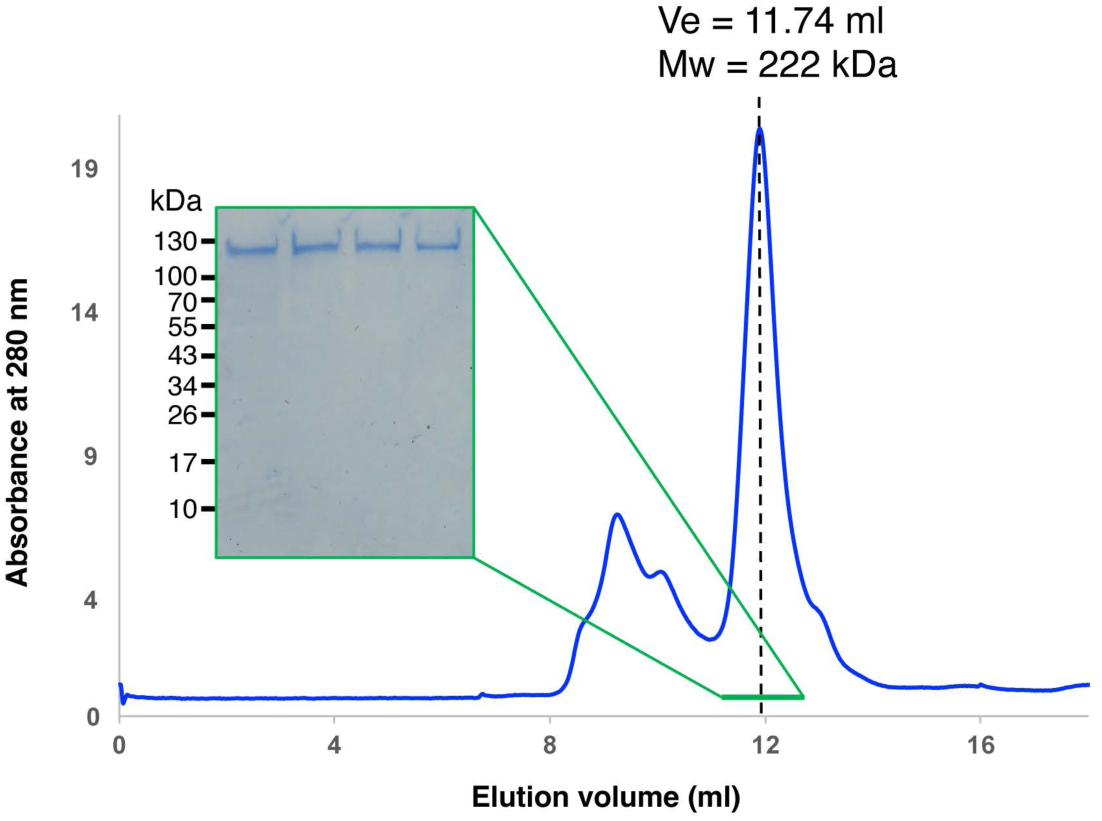
NbNRC2 elutes as a dimer in analytical gel filtration assays. Gel filtration assays with NbNRC2. C-terminally 6xHis-3xFLAG tagged NbNRC2 was expressed and purified from leaves of *nrc2/3/4* KO *N. benthamiana*. Purified protein was run on an S200 10/300 analytical column, and the estimated molecular weight of the NRC2 peak was calculated based on comparison of the elution volume to a protein standard curve ran on the same column. Elution volume (Ve) and molecular weight (Mw) in kDa are shown above the peak. A range of fractions below the peak, marked in green, were ran on SDS–PAGE and visualized using Coomassie Brilliant Blue staining (green panel). Approximate molecular weights (kDa) of the proteins are shown on the left of the green panel.

**Fig. S2:**
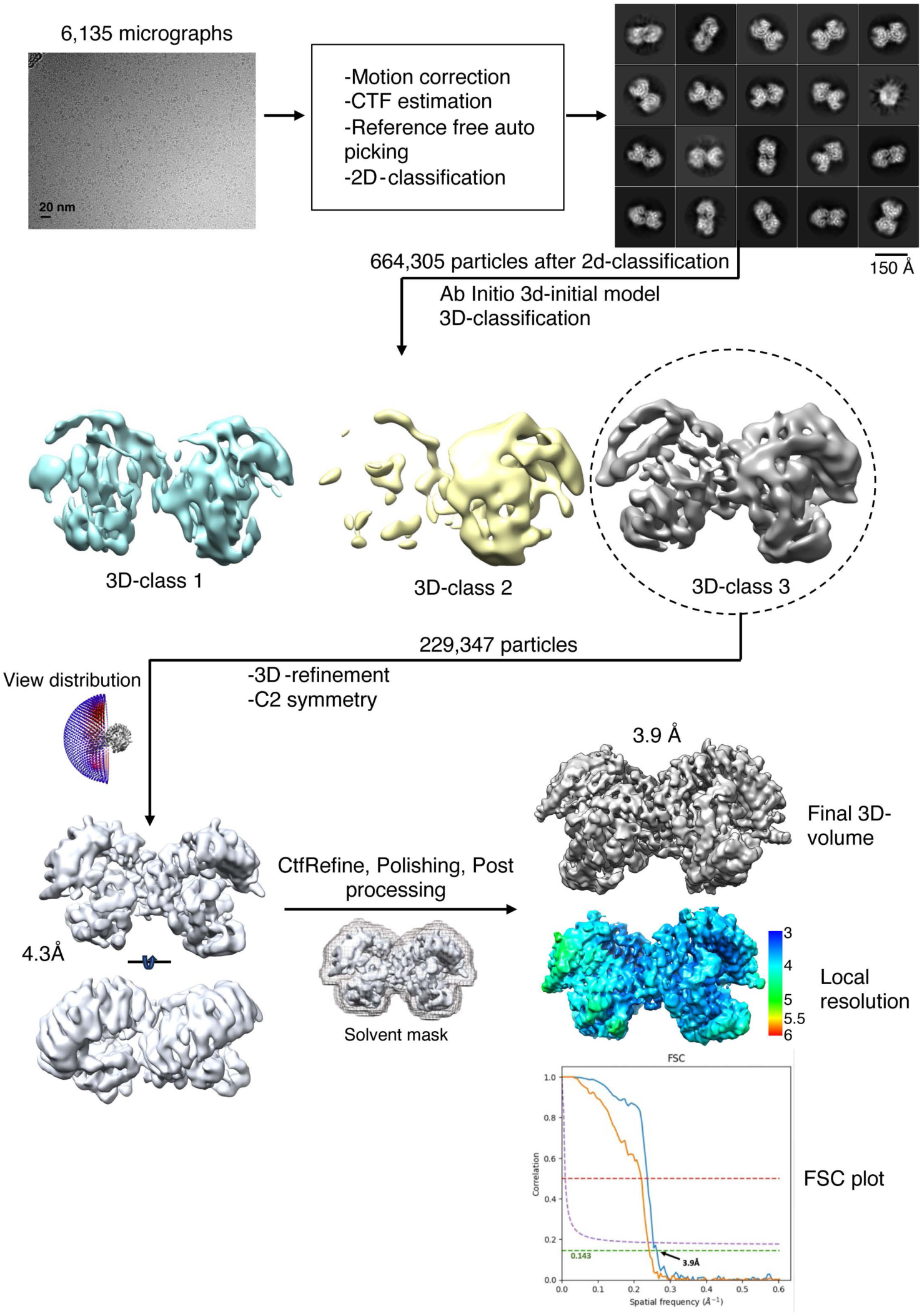
Cryo-EM image processing and 3D-reconstruction of NRC2 homodimers. Schematic representation of Cryo-EM data processing and 3D-reconstruction of resting state NRC2 homodimer. Detailed information on the data collection and processing as well as model building can be found in the pertinent Methods section and in **Table S1**.

**Fig. S3:**
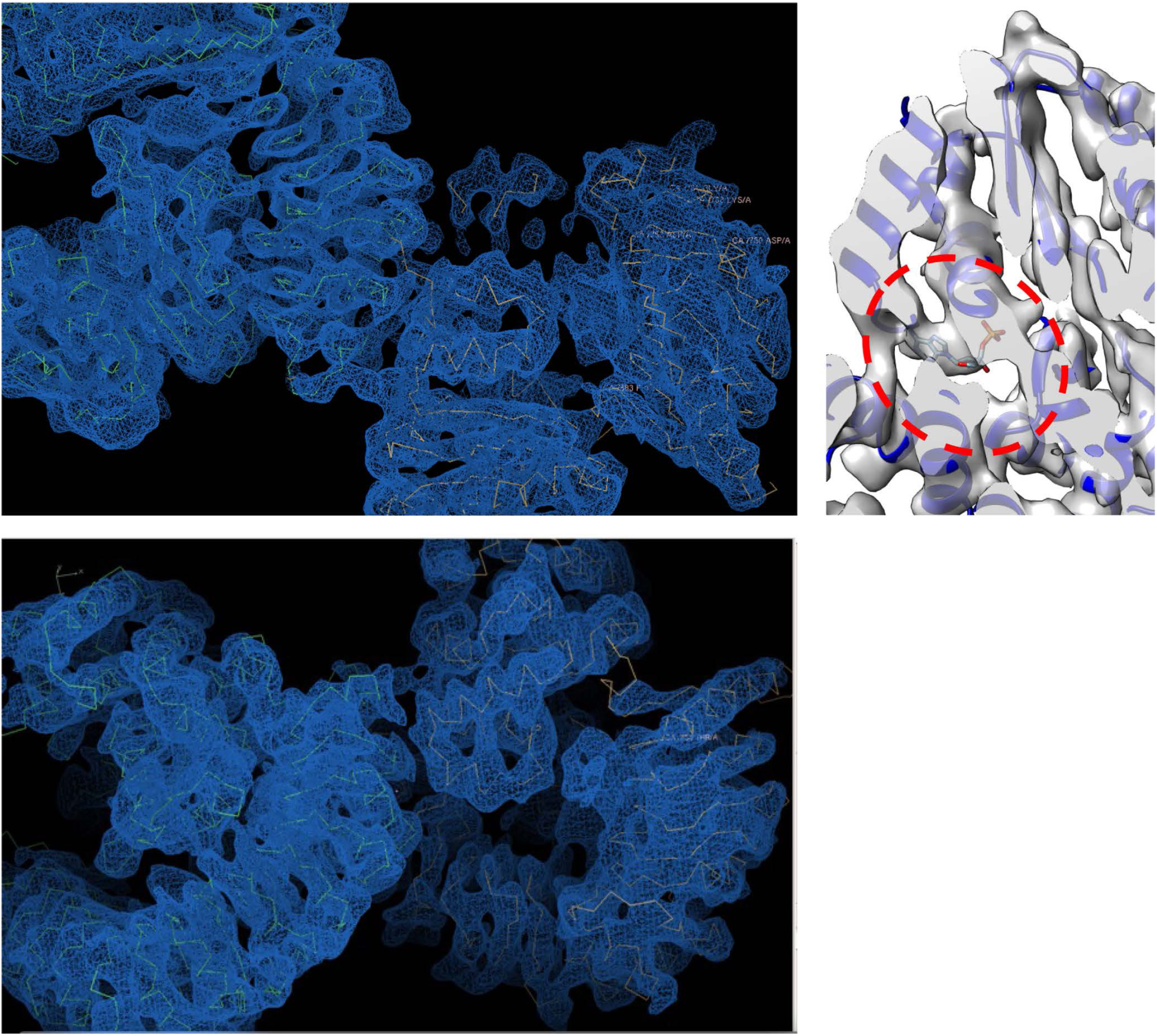
Representative map sections of NRC2 homodimer showing the overall fit of the atomic model of NRC2 into the Cryo-EM density map. The extra density at NB domain which accounts for ADP is circled.

**Fig. S4:**
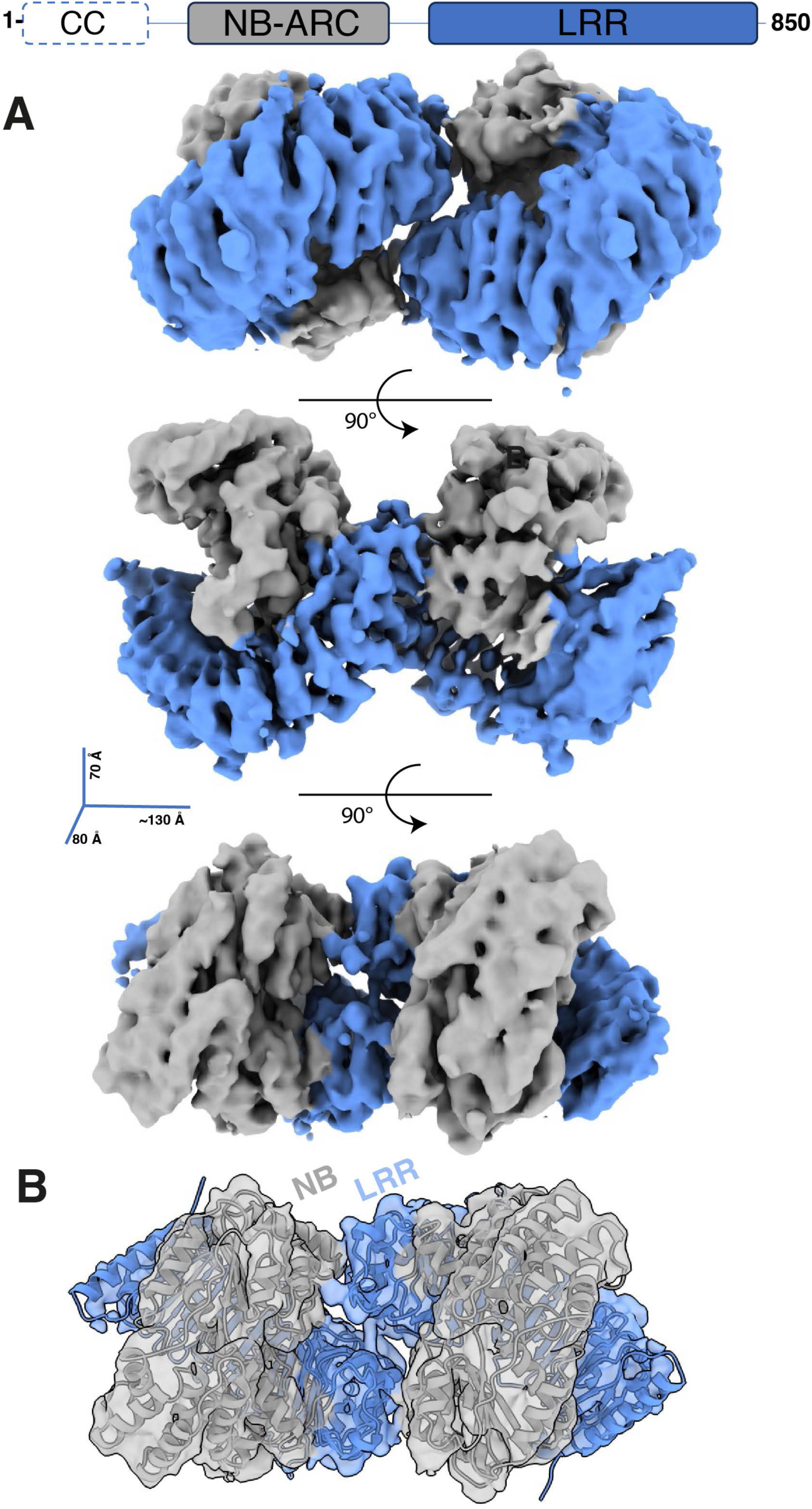
The helper NLR NRC2 forms a homodimer *in planta*. (**A**) Overall molecular architecture and organization of resting state NRC2 homodimer. Schematic of NbNRC2 domain architecture is shown above. (B) Fit of NRC2 AlphaFold2 model into the Cryo-EM density map.

**Fig. S5:**
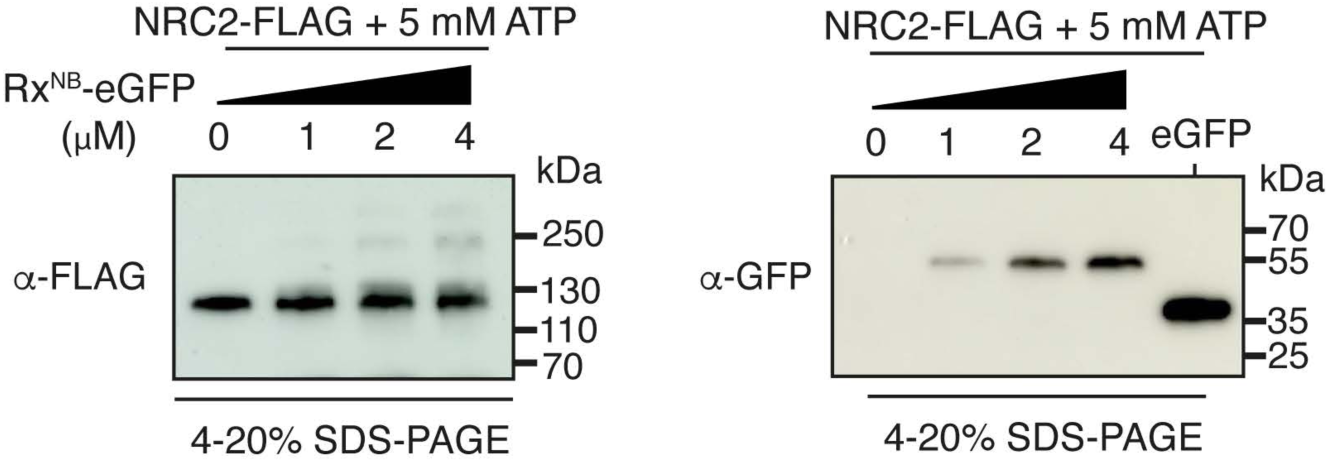
SDS-PAGE assays accompanying BN-PAGE assays in. Fig. 2B. Affinity-purified proteins were run on SDS-PAGE assays and immunoblotted with the appropriate antisera labelled on the left. Approximate molecular weights (kDa) of the proteins are shown on the right.

**Fig. S6:**
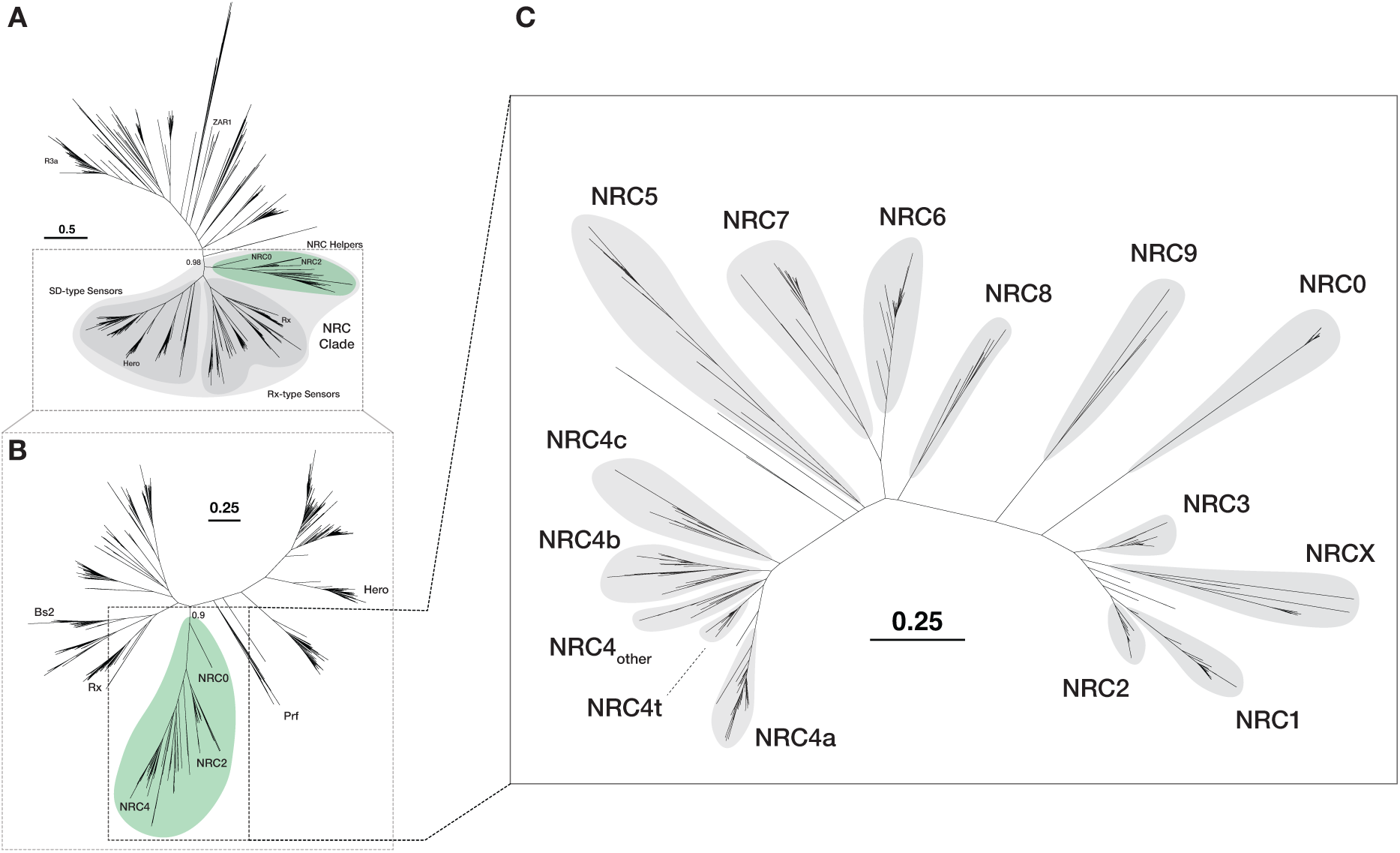
NRC sequences form a well-supported monophyletic clade within the CC-NLR and NRC superclade phylogenetic trees. (**A**) NRC superclade including NRC helpers, Rx-type sensors, and SD-type sensors form a distinct, well-supported clade in a phylogenetic tree of 19,760 CC-NLRs from the Solanaceae family. (**Data S6**, **Data S7**) (**B**) NRCs are monophyletic and part of the same well-supported clade inside the NRC superclade phylogenetic tree. (**Data S8**) (**C**) NRC sequences are divided into 15 distinct phylogenetic subclades based on the presence of reference NRC sequences (**Data S9**, **Data S10**). Numbers next to the major branches indicate bootstrap values.

**Fig. S7:**
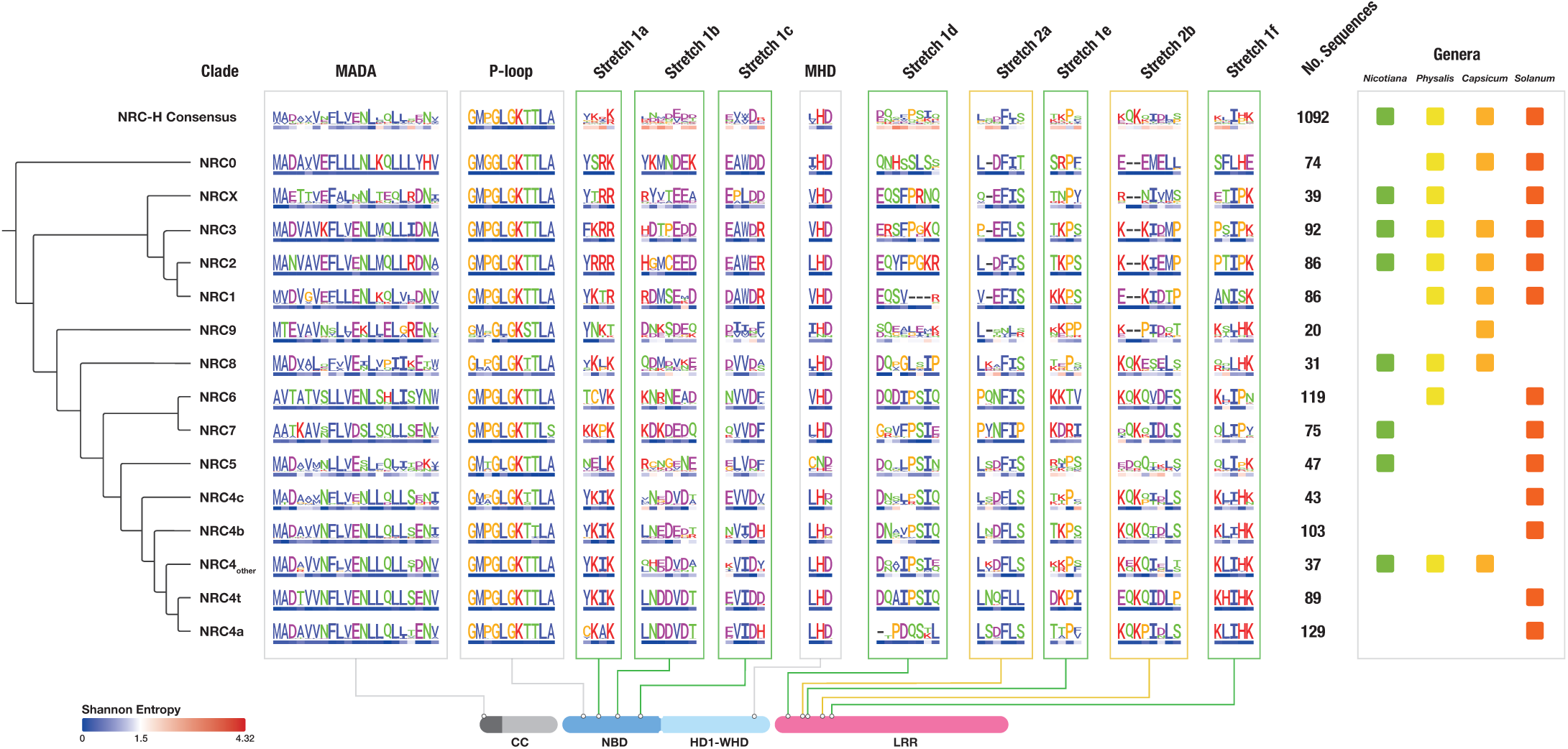
Sequence motif and Shannon entropy plots for 1,092 NRC sequences retrieved from 123 genome assemblies from 4 genera in the Solanaceae family. Sequence motif and Shannon entropy plots for the MADA, P-loop, MHD motifs, and eight dimerization stretches of 1,092 NRC sequences grouped into NRC clades based on phylogenetic relationships. The number of sequences and the distribution across the 4 genera within each clade are indicated on the right. (**Data S3**, **Data S4**).

**Fig. S8:**
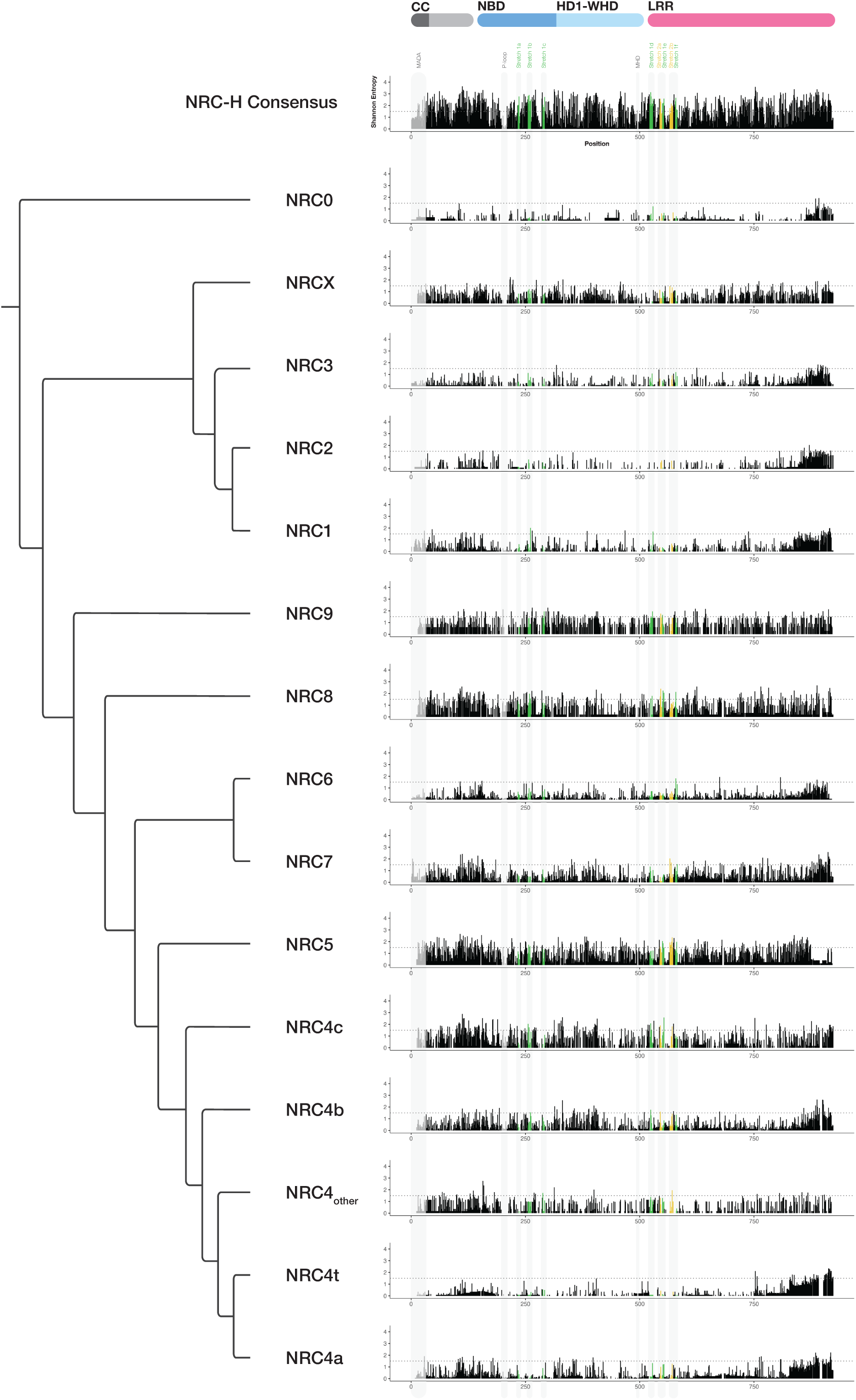
Shannon entropy plots of full-length proteins calculated across all 1,092 NRC sequences and within each individual NRC clade. The dimer stretches 1 and 2 are colored by green and yellow, respectively. MADA, P-loop and MHD motif are colored by grey. Shannon entropy score of 1.5, indicated by the dotted line, was established as a threshold to distinguish between highly variable (>1.5 bits) and non-highly variable (<1.5 bits) amino acids, as established previously (31).

**Table S1:** Cryo-EM data collection, refinement, and validation statistics.

| **Data collection and processing** | **Resting state NRC2**<br>**PDB: 8RFH - EMDB: 19121** |
| --- | --- |
| Microscope | Titan Krios |
| Voltage(KeV) | 300 |
| Detector | Gatan K3 |
| Magnification | 105,000 |
| Calibrated pixel size (Å) | 0.828 |
| Exposure time (s) | 2.6 |
| Frames per exposure (e<sup>-</sup>/Å<sup>2</sup>) | 50 |
| Total electron exposure | 50 |
| Automation software | EPU |
| Defocus range (µM) | 1.5 to 2.7 |
| Number of micrographs used | 6135 |
| Total refined particles (no.) | 229,347 |
| Symmetry imposed | C2 |
| Map sharpening B-factor (Å<sup>2</sup>) | -100 |
| Unmasked Resolution at 0.5/0.143 FSC (Å) | 4.1/4.5 |
| Masked resolution at 0.143/0.5 FSC (Å) | 3.9/4.2 |
| **Model refinement and validation statistics** |  |
| PDB composition | 2 Protein chains |
| Amino acids | 1469 |
| Ligand | ADP |
| RMSD bonds (Å) | 0.002 |
| RMSD angles (°) | 0.773 |
| B-factor | 176 |
| Ramachandran |  |
| Favoured (%) | 97.51 |
| Allowed (%) | 2.49 |
| Outliers (%) | 0.0 |
| Rotamer outliers (%) | 0.30 |
| Clash score | 15.89 |
| C-beta outliers (%) | 0.0 |
| CaBLAM outliers (%) | 0.97 |
| CC (mask) | 0.75 |
| CC (volume) | 0.732 |
| Molprobity score | 1.8 |
| EMRinger | 0.87 |

**Table S2:** Interface residues at 6 Å cut-off distance and their definition as amino acid stretches.

|  |  |
| --- | --- |
| **Interfaces 1 and 3** | <u>NB domain region:</u><br>Y217,R220,H238,D239,M240,C241,E242,E243,D244. D270 and R274.<br><u>LRR domain region:</u><br>R513,K512,G511,S536,K563,P559,F509,Y508,E506,K534,T560,S532,P535,T533 |
| **Interface 2** | <u>LRR region:</u><br>Q507,S532,E529, L528,P554,M553,E552,K549 |
| **Amino acid stretches involved in interfaces 1 and 3** | • Stretch1a:217-220<br>• Stretch1b: 238-244<br>• Stretch1c: 270-274<br>• Stretch1d: 506-513<br>• Stretch 1e: 533-536<br>• Stretch 1f: 559-563 |
| **Amino acid stretches involved in interface 2** | • Stretch2a: 528-532<br>• Stretch 2b: 549-554 |

**Movie S1: 3D view of NRC2 homodimer reconstructed from Cryo-EM micrographs (as separate file).** NB-ARC domain is shown in gray. LRR domain is shown in blue.

**Data S1: Sequence alignment of NbNRC2, NbNRC3, NbNRC4, and NbZAR1 (as separate file).**

**Data S2: Genome metadata used in the phylogenetic analyses (as separate file). Data S3: Curated 1,092 NRC sequences in FASTA format (as separate file).**

**Data S4: Entropy values for 1,092 NRC sequences and each NRC clade for the analysed dimerization stretches, sequences motifs, and the full-length proteins (as separate file).**

**Data S5: Frequencies of each amino acid within the NRC2, NRC3, NRC4other, NRC4a, and NRC4b clades compared to other NRC clades (as separate file).**

**Data S6: List of CC-type NLR sequences and associated metadata (as separate file).**

**Data S7: Extracted NB-ARC (NBD and HD1-WHD) domain sequences of CC-type and reference NLRs (as separate file).**

**Data S8: NB-ARC (NBD and HD1-WHD) domain sequences of NRC superclade and reference NLRs (as separate file).**

**Data S9: NB-ARC (NBD and HD1-WHD) domain sequences of curated NRC sequences (as separate file).**

**Data S10: NB-ARC (NBD and HD1-WHD) domain sequences of reference NRC sequences from *S. lycopersicum*, *S. tuberosum*, *N. benthamiana*, *C. annuum*, and RefPlantNLR (as separate file)**.

